# Ontogeny of risk assessment and escape-hatching performance by red-eyed treefrog embryos in two threat contexts

**DOI:** 10.1101/2022.05.09.491251

**Authors:** Brandon A. Güell, Julie Jung, Adeline Almanzar, Juliana Cuccaro-Díaz, Karen M. Warkentin

**Author notes:** Correspondence: K. M. Warkentin, Department of Biology, Boston University, 5 Cummington Mall, Boston, MA 02215, U.S.A.

## Abstract

Arboreal embryos of red-eyed treefrogs, *Agalychnis callidryas*, hatch prematurely in response to hypoxia when flooded and mechanosensory cues (MC) in snake attacks, but hatching later improves tadpole survival. We studied ontogenetic changes in risk assessment and hatching performance of embryos in response to flooding and physical disturbance. We hypothesized that risk assessment decreases as hatchling survival improves and hatching performance increases as embryos develop. Because snakes eat faster than embryos asphyxiate, we hypothesized that embryos decide to hatch sooner and hatch faster in response to MC. We video-recorded individual embryos hatching in response to each cue type, then compared the incidence and timing of a series of events and behaviors from cue onset to complete hatching across ages and stimuli. Latency from cue to hatching decreased developmentally in both contexts and was shorter with MC, but the elements contributing to those changes differed. Hypoxia-assessment involved position changes, which decreased developmentally along with assessment time. MC-assessment was passive, faster, and also decreased with age. The first stages of hatching, membrane rupture and head emergence, were surprisingly age-independent but faster with MC, congruent with greater effort under more immediate risk. In contrast, body emergence and compression showed ontogenetic improvement consistent with morphological constraints but no cue effect. Both appropriate timing and effective performance of hatching are necessary for continued development. Different stages of the process vary with development and environmental context, suggesting combinations of adaptive context- and stage-dependent behavior, cue-related constraints on information acquisition, and ontogenetic constraints on elements of performance.

**Summary statement:** Development reduces risk assessment and improves hatching performance by red-eyed treefrog embryos in both flooding and simulated attacks. Predation cues elicit faster decisions and hatching, congruent with more immediate risk.

## INTRODUCTION

Adaptive animal behavior relies on effective use of information and performance of actions (Dall et al., 2005). As animals develop, their sensory (Gervais et al., 2021; Romagny et al., 2012) and motor capabilities (Bate, 1999) change, affecting how they perceive cues and physically interact with their environment (Danchin et al., 2004; Wiedenmayer, 2009). Moreover, in order to respond with adaptive behaviors, animals must also balance the value of gathering information with its cost (e.g., sampling time, energy expenditure) (Bradbury and Vehrencamp, 1998; Dall et al., 2005; Warkentin and Caldwell, 2009). However, development changes the costs and benefits of performing specific behaviors in a given context, and therefore also alters which responses are best at different points in ontogeny (Wiedenmayer, 2009). Here, we test how development affects the decision-making and performance processes of an essential animal behavior in response to two different threat cues, associated with different costs of sampling and latencies to mortality.

For oviparous animals, hatching is an essential and often behavioral process that developing embryos must perform. It typically requires the use of specific mechanisms to rupture the egg capsule and behaviors to exit from it (Bles, 1906; Cohen et al., 2018; Cohen et al., 2016; Oppenheim, 1972; Yamagami, 1981; Yamagami, 1988). For many species this can be a physically demanding process since developing embryos are often enclosed in multiple layers of membranes, jelly, shells, etc. that provide protection (Altig and McDiarmid, 2007; Dumont and Brummett, 1985). Hatching may be particularly challenging for younger, less developed embryos in species that have long plastic hatching periods (Warkentin, 2011). For instance, if hatching performance traits develop gradually, as a result of developing bodies and physical abilities, embryos may pass through an initial period of marginal hatching competence. In such cases, if partial hatching by less developed embryos compromises the protective functions of egg capsules, then hatching complications or failure may themselves cause embryo mortality, selecting against hatching attempts and increasing the premium on risk assessment and decision accuracy.

Adaptively plastic timing of hatching, cued by environmental conditions, is phylogenetically widespread (Warkentin, 2011). Environmentally cued hatching allows embryos to navigate fitness trade-offs to determine the optimal time to hatch in response to a variety of stimuli. Embryos often use prolonged or repeated sampling to inform their hatching decisions and balance the costs of missed cues and false alarms (Warkentin and Caldwell, 2009). However, as sampling increases, so does its cost. Assessment costs can be crucial for embryos sampling cues associated with a source of mortality (e.g., egg-predator cues), since increasing lag time before hatching in these contexts increases the likelihood of death (Warkentin and Caldwell, 2009).

Embryos should therefore adjust sampling of cues from different sources based on how the value and cost of information accrue. Development can also affect how embryos assess risk-cues if the costs of sampling their environment, the hatching process, or entry into their next life stage change ontogenetically. Here, we use two different cues—hypoxia and physical disturbance— that indicate common threats to terrestrial eggs (flooding and predation) to assess how development changes the risk assessment, decision-making and hatching process in a well-studied example of environmentally cued hatching.

The terrestrial eggs of red-eyed treefrogs, *Agalychnis callidryas*, offer an excellent system for direct observations of embryo development and behavior in several induced-hatching contexts. Undisturbed embryos typically hatch at age 6–7 days but use multiple sensory modalities to assess risk of flooding and predation, hatching in response to hypoxia and mechanosensory cues (Jung et al., 2019; Jung et al., 2020; Warkentin, 2002; Warkentin, 2005). Hypoxia-cued hatching begins at age 3 days and mechanosensory-cued hatching begins at age 4 days (Warkentin et al., 2017). However, neither hypoxia nor physical disturbance consistently indicate real threats to eggs. In particular, due to spatial gradients of oxygen within eggs, embryos often experience transient hypoxia due to their orientation within the egg, which they solve by changing position (Rogge and Warkentin, 2008). Embryos may also experience more persistent hypoxia if pond levels rise to submerge clutches, or if individual eggs or entire clutches fall into the water.

Similarly, rainstorms produce intense vibrations with properties that overlap those in predator attacks, yet they pose no threat to eggs (Warkentin, 2005). This overlap in cue properties from benign and threatening sources of stimuli creates a discrimination challenge for embryos and necessitates adequate sampling to avoid false alarms and make informed decisions of whether and when to hatch. However, the cost of false alarms decreases developmentally; older hatchlings are larger, more developed, more behaviorally competent, and suffer lower mortality in the water, particularly with aquatic predators (Gibbons and George, 2013; Touchon et al., 2013; Warkentin, 1995; Warkentin, 1999a; Willink et al., 2014). In contrast, the ultimate cost of missed cues—death by asphyxiation or consumption by a predator—does not change across development (Warkentin and Caldwell, 2009).

Cued hatching in *A. callidryas* is mediated by rapid, localized enzyme release by two types of hatching gland cells (HGC) (Cohen et al., 2019). Early HGC appear at 3 days, begin to regress at 4 days, and enable the earliest cued hatching, while late HGC appear at 4 days, gradually increase in abundance, and mediate most hatching events (Cohen et al., 2019). Before membrane rupture and hatching, most embryos exhibit shaking behaviors associated with hatching enzyme release, indicating the onset of the hatching process (Cohen et al., 2016). Typically, embryos maintain their snout at the rupture site and use thrashing movements to propel themselves through the hole (Cohen et al., 2016). Developmental stage at hatching could affect the process or performance of hatching if using late HGC, or more HGC, accelerates the process of membrane rupture or increases the size of the hole produced, thereby facilitating exit from the capsule. Moreover, as embryos develop they also increase in total size, their axial musculature increases, and they become more streamlined as their heads grow and bulbous yolk sac transforms into gut coils (Warkentin, 1999b). These gross morphological changes may also facilitate their exit through a small membrane rupture and improve hatching performance. Thus, we hypothesize that hatching performance improves developmentally in response to both hypoxia and physical disturbance cues, resulting in older embryos hatching faster either overall or through specific periods within the process. Moreover, because the risk of mortality is more immediate and accrues more quickly in predator attacks than during flooding (i.e., rapid consumption vs. gradual asphyxiation) (Warkentin and Caldwell, 2009; Warkentin et al., 2007), embryos cued by physical disturbance may exhibit hatching performance closer to their maximum capacity. If so, we predict that the process of hatching—from initiation to exit from the egg—is faster in response to physical disturbance cues and that performance varies more in hypoxia-cued hatching than in mechanosensory-cued hatching.

*Agalychnis callidryas* embryos have distinct and measurable periods of cue sampling before deciding to hatch. For instance, when deprived of oxygen, either briefly when misoriented within their egg or for prolonged periods when flooded, embryos’ first response is to reorient themselves within their eggs, changing position repeatedly in attempt to return their gills to air-exposed parts of the egg (Rogge and Warkentin, 2008; Warkentin et al., 2017) (Movie 1). These position changes are distinct from the movements associated with hatching and are therefore useful indicators of oxygen sampling. However, sampling oxygen must come at some metabolic and sampling time cost, since embryos actively move their bodies to reposition their gills, stirring the perivitelline fluid, and must wait for oxygen level to stabilize before they can accurately assess it at each new position. Moreover, embryos pay an increasing developmental cost the longer they remain in the egg under hypoxic conditions (Snyder et al., 2018; Vasquez et al., 2016). In contrast, sampling vibrations and tactile cues in physical disturbance is passive, and information accrues more rapidly (Caldwell et al., 2009; Warkentin and Caldwell, 2009), although embryos risk the greater and more immediate threat of being eaten while sampling this type of information (Warkentin and Caldwell, 2009). Since hatchling mortality decreases with development, younger embryos should spend more time sampling both cue types before deciding to hatch, and younger embryos should also sample more positions when flooded. Sampling periods of age-matched embryos should be shorter in response to physical disturbance cues because mortality accrues faster in predator attacks.

Previous work has used latency from stimulus onset to hatching to estimate cue sampling (Jung et al., 2019; Jung et al., 2020; Warkentin et al., 2017; Warkentin et al., 2019) and found that latency to hatch in response to vibration playbacks decreases from age 5 to 6 days (Warkentin et al., 2019). However, this latency period includes the time required to rupture and exit the egg in addition to the risk-assessment and decision-making processes. Thus, latency from stimulus onset to hatching initiation may be a more accurate measure of cue sampling. Moreover, to understand if changes in hatching behavior occur evenly across development or if specific changes are concentrated in shorter periods, associated with specific developmental changes, it is necessary to measure ontogenetic changes in both hatching performance and risk assessment across the full period of hatching competence. Thus, we video-recorded individual *A. callidryas* embryos at ages 3–6 days hatching in response to hypoxia (flooding) and physical disturbance (jiggling) cues and compared the occurrence and timing of specific behaviors and periods within the risk-assessment and hatching processes across ages.

## MATERIALS AND METHODS

### Egg clutch collection and care

We collected young (0–3 days old) *A. callidryas* egg clutches on leaves from the Experimental Pond in Gamboa, Panama (9°7 ′15 ″N, 79°42 ′14 ″W), attached the leaves to plastic support cards, and placed them in cups over aged, dechlorinated tap water to catch hatchlings. Clutches were maintained in an open-air, ambient temperature and humidity laboratory at the Smithsonian Tropical Research Institute (STRI) in large plastic bins, with screen windows in the lids to allow air flow and misted frequently with rainwater to maintain hydration. All embryos used were morphologically normal, in developmental synchrony with siblings in their clutch, and in intact, turgid eggs at the start of testing. Most eggs are laid between 22:00 h and 02:00 h, so we assigned embryo ages starting from midnight of their oviposition night (Warkentin, 2002; Warkentin et al., 2005). We returned all hatched tadpoles to the Experimental Pond after experiments. All research was conducted under STRI IACUC protocol 2014-0601-2017 and Boston University IACUC protocol 14-008 and permits from the Panamanian Ministry of the Environment (SC/A-15-14, SE/A-46-15, SE/A-59-16).

### Hypoxia-cued hatching

To assess developmental changes in hypoxia-cued hatching, we video-recorded flooded embryos hatching at four ages, from 3–6 days. At age 3 days, before embryos hatch in response to physical disturbance cues (Warkentin et al., 2017), we placed individual eggs in custom-made glass egg-cups (Fig. 1, Fiamma Glass, Waltham, MA). The cups were designed to fit eggs closely (interior diameter ca. 5 mm, depth 3–4 mm) and mounted on glass bases with their opening vertically oriented (Fig. 1A). Eggs in cups were thus exposed to air on one side (henceforth the “front”), but the glass — rather than sibling eggs or leaf — blocked gas exchange through about half of their surface, providing a standardized, yet naturalistic, oxygen environment (Rogge and Warkentin, 2008; Warkentin, 2002; Warkentin et al., 2005) (Movie 1). Egg-cups allowed us to move eggs without touching them, facilitating manipulation of older embryos, and improved the visibility of behaviors. Based on embryos’ external morphology, rearing in cups did not alter development. We placed eggs-in-cups in a plastic container with a screened window in the lid and misted the eggs and interior of the container frequently with rainwater to maintain hydration.

**Figure 1.**
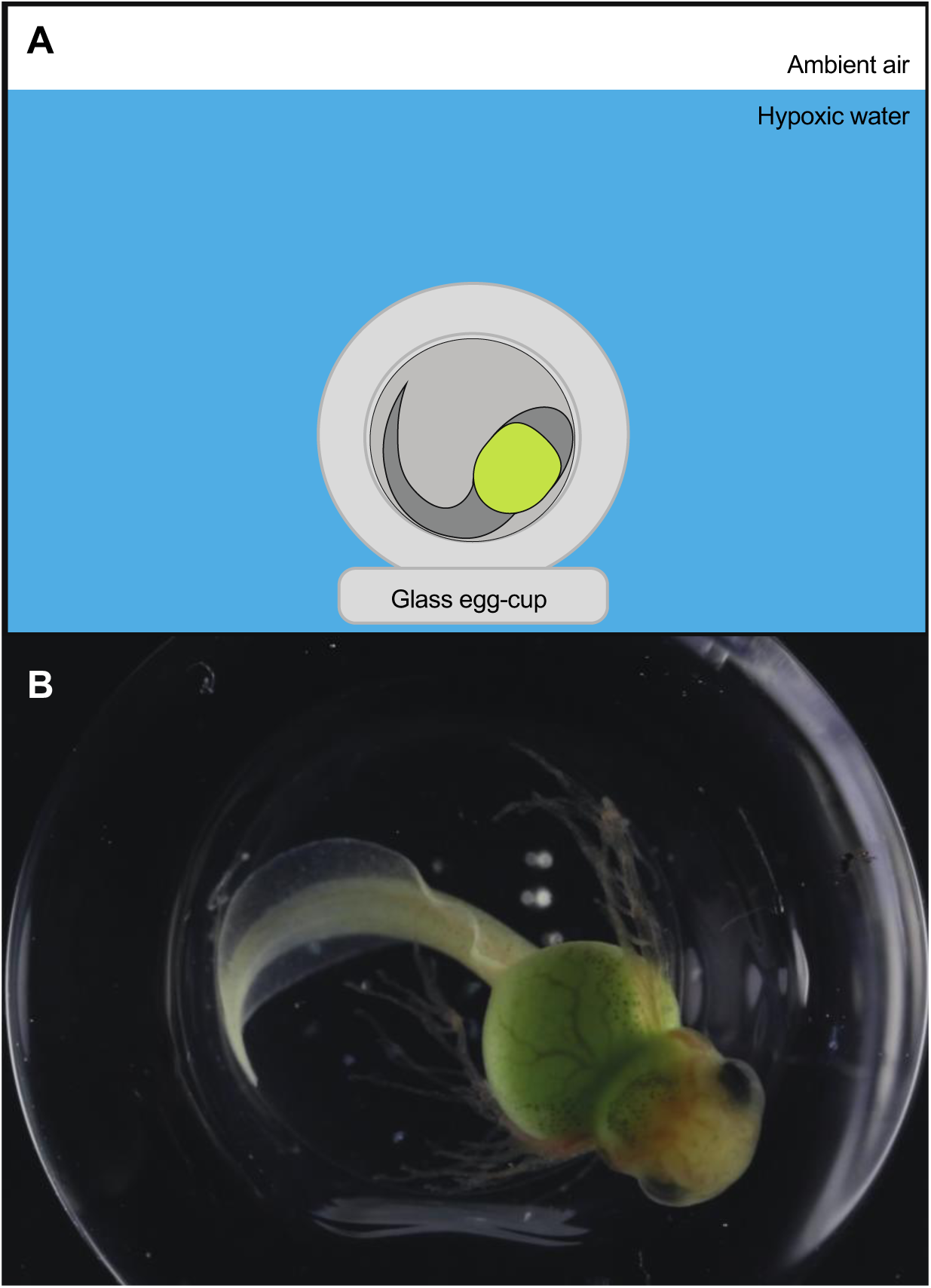
Methods for video-recording hypoxia-cued hatching of individual *Agalychnis callidryas* embryos. (A) Focal embryo in a glass egg-cup submerged in hypoxic water in a glass-fronted Plexiglas tank. Illustration is not to scale. (B) Cropped camera view of an *A. callidryas* embryo at age 3 days in the process of hatching. Body compression is evident (see also Movie 3).

We tested embryos at ages 3–5 days during June and July 2014, and embryos at ages 5–6 days during June and July 2015. We began testing embryos at age 3 days at 18:00 h, when all sibships were hatching-competent (Warkentin et al., 2017), and we tested embryos from ages 4–6 days between 08:00 h and 16:00 h. In 2014, we tested embryos in a small aquarium constructed from a Plexiglas tube (5 cm diameter, 6.8 cm high) with a glass front inserted creating an optically clear area (3.5 cm wide x 4 cm high) for viewing and recording. To begin a test we moved an egg-in-cup into the tank (Fig. 1), waited 5 minutes to ensure that the setup process had not stimulated hatching, then began recording. We recorded videos at 30 frames per second using a Canon EOS 5D Mark III camera and MP-E 65 mm macro lens, with the egg illuminated from both sides using two LED lights. We gently flooded the aquarium with hypoxic water to submerge eggs (Movie 1). To make hypoxic water, we boiled tap water for at least 10 minutes, sealed it in glass-stoppered BOD bottles without air bubbles, and allowed it to cool. Water was used within 30 min of opening the bottle (15±1.3% air saturated at opening, 21±3% air saturated at 30 min, *N*=10 bottles, mean±s.d. here and in text throughout). We recorded video until the embryo hatched, then moved hatchlings to air-saturated water. We photographed all hatchlings in dorsal view with a ruler, then measured hatchling size from photographs using ImageJ (Schneider et al., 2012). The few embryos at age 3 days that failed to hatch were removed from the test chamber after 40 minutes and either returned to air, in their egg-cups, or manually decapsulated and placed into air-saturated water.

Moving eggs-in-cups into the video-recording tank at age 6 days induced hatching. Therefore, in 2015, we set up eggs-in-cups in sets of four in larger video-recording tanks (width, depth, height:10.3 × 4.9 × 8 cm) at age 3 or 4 days and left them undisturbed until testing. We misted the eggs-in-cups and interior of tanks with rainwater frequently and covered the tanks with dampened mesh to maintain eggs hydrated until testing. At testing, we focused the camera on a single intact egg, flooded the tank as above, and recorded the behavior and hatching of the embryo. We used these methods for five additional embryos tested at age 5 days and all embryos tested at age 6 days. Since the two setup methods did not alter the hatching behavior or performance of embryos at age 5 days, we pooled the data for analysis.

### Mechanosensory-cued hatching

To assess developmental changes in mechanosensory-cued hatching, we manually jiggled eggs, following methods from Warkentin et al. (2017), and video-recorded hatching at three ages, 4–6 days. Jiggling does not induce hatching of embryos at age 3 days, due to insufficient development of mechanosensory systems (Jung et al., 2019; Jung et al., 2020). For each test, we removed an individual egg from its clutch, placed it in a small hexagonal weigh boat with a drop of water, and manually jiggled the egg using a moistened blunt metal probe, alternating 15 s of stimulation with 15 s of rest for 5 minutes or until the egg hatched (see Video S1 in Warkentin et al., 2017).

We conducted mechanosensory-cued hatching tests during June–August 2016 between 12:00 h and 18:00 h. For embryos at ages 4 and 5 days, and for most embryos at age 6 days (*N*=6 of 11), we waited at least 5 minutes after transferring the egg to the testing dish before starting to record video and jiggle the egg (4 d: 6.86±8.24 min, 5 d: 6.44±1.19 min, 6 d: 5.59±3.85 min). Since moving eggs from clutches to their weigh boats at age 6 days often induced hatching, 4 individuals tested at age 6 days were acclimated for less than 5 minutes (2 individuals for 1.5 min, 2 individuals for 1.75 min). For 1 individual that hatched immediately at set up, we included egg transportation and setup as physical disturbances which induced the recorded hatching behavior. In these cases, we started the recording before picking the egg off the clutch and narrated the timing onto the audio track. All hatchlings were photographed and measured as above.

### Video analysis

We recorded a total of 148 individual embryos hatching in response to hypoxia cues, including up to 2 individuals per age per clutch and up to 3 ages per clutch. Of these, we analyzed 45 recordings from 32 clutches (*N*=10, 10, 15, and 10 individuals from 10, 10, 12, and 10 clutches at ages 3–6 days, respectively) that met the following criteria: i) the embryo had an undisturbed 5-minute acclimation period; ii) the flooding process did not tip or shift the glass cup from its original position; iii) the initial membrane rupture was made in the front, exposed portion of the egg, not against the glass; iv) the embryo exited through the initial membrane rupture; and v) the embryo completely exited from its egg capsule without obstruction by the glass cup. We recorded a total of 100 individual embryos hatching in response to physical disturbance cues (up to 2 individuals per age per clutch and up to 2 ages per clutch) and analyzed a total of 45 recordings from 39 clutches (*N*=18, 16, and 11 individuals from 16, 16, and 10 clutches at ages 4–6 days, respectively) that met the following criteria: i) physical disturbance did not continue after the embryo began performing hatching behavior; ii) the embryo exited through the initial membrane rupture; and iii) the timing of events was accurately measurable from the video recording and narration.

From the videos, we quantified the occurrence and timing of a series of events and behaviors within the hatching process, some of which are described elsewhere (Cohen et al., 2016), and compared them across ages and stimuli. We defined cue-sampling duration as the period from stimulus onset (complete submergence in hypoxic water or start of egg manipulation in physical disturbance) to when behaviors indicating the onset of hatching began. We also counted the number of times embryos changed position within the egg throughout this cue-sampling period. As indicators of the initiation of hatching, we used four behaviors that are often expressed shortly before membrane rupture: shaking, mouth gaping, jerks, and buccal cavity compressions. Shaking refers to axial muscle contractions that generate low-amplitude lateral movements (Cohen et al., 2016). Mouth gaping resembles buccal pumping but with larger amplitude movements and extended duration of the gape (Cohen et al., 2016). Jerks consist of single axial muscle contractions that are stronger than an individual contraction within shaking behavior (Movie 2). Buccal cavity compressions cause a brief change in the shape of the snout, without gaping open the mouth or jerking the body (Movie 2). Since hatching enzyme release has been experimentally shown to occur during shaking (Cohen et al., 2016), we used the onset of shaking to estimate hatching initiation if embryos exhibited this behavior. If shaking did not occur, we used one of the alternative behavioral indicators that embryos performed prior to membrane rupture, with their snout positioned at the location of the subsequent rupture, to estimate the beginning of hatching (*N*=1, 1, 2, and 4 embryos at ages 3–6 days, in hypoxia trials only). We were unable to clearly assess the timing of any behavioral indicator in three jiggling trials. We assessed the timing of key events within the hatching process with 1/30 s (single frame) accuracy and used these to calculate the following periods: hatching initiation to membrane rupture, membrane rupture to head emergence, head emergence to body emergence (excluding the tail), hatching initiation to complete exit from the egg, and start of thrashing to complete exit from the egg. Thrashing motions are performed by most embryos before and during their exit from the capsule and consist of high-amplitude body undulations that travel from snout to tail (Cohen et al., 2016). In two cases, embryos successfully hatched but rested for an extended period with their tail tip within the capsule; we considered these to have exited the egg when their body emerged from the membrane. Since we recorded video at 30 frames per second, we could not accurately measure shorter durations; for analysis we assigned durations of 1/30 s to all processes completed in ≤1/30 s (*N*=6).

### Statistics

We used linear and generalized linear mixed models (‘lme4’ package; Bates et al., 2015) with clutch as a random effect, followed by likelihood ratio tests of nested models to determine the main effects of age (coded as ordinal), cue type, and their interaction. When age × cue type interaction effects were significant in our original models, we ran additional, independent mixed models on data within each cue type, followed by likelihood ratio tests for age effects. When overall age effects were significant within cue types, we used Tukey *post hoc* tests (‘multcomp’ package; Hothorn et al., 2008) to determine differences between specific ages. To test for effects of cue type on variation in hatching latency and elements of hatching performance, within ages, we used Fligner-Killeen tests of homogeneity of variances. We used linear mixed models on natural-log-transformed data when original data did not fit gaussian, gamma, or binomial error distributions. We performed *t*-tests and Wilcoxon signed-rank tests to determine if sampling durations and number of position changes were statistically different from zero at age 6 days. We used a generalized linear model to test if time to thrashing onset predicts time to head emergence, measuring both from membrane rupture. All statistical tests were performed in the R statistical environment (R Core Development Team, 2019) using RStudio (version 1.0.143).

## RESULTS

### Hatchling size

As expected, age was the strongest predictor of hatchling size (LMM, main effect of age: χ^2^=89.664, *P*<2.2e-16) regardless of whether we included hatchlings at age 3 days from hypoxia tests (Fig. 2). Hatchling size increased developmentally in both hypoxia (χ^2^=59.972, *P*=5.96e-13) and physical disturbance tests (χ^2^=12.336, *P*=0.002, Fig. 2).

**Figure 2.**
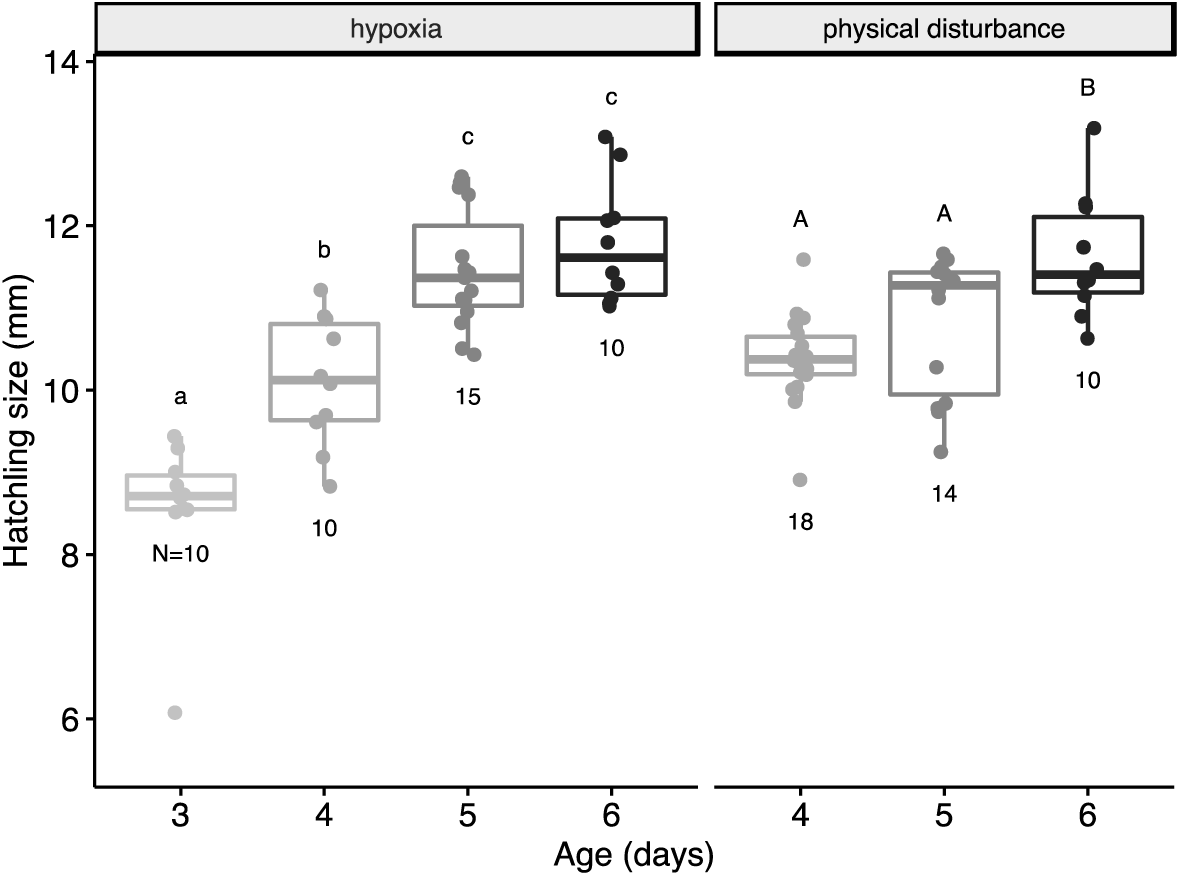
Total lengths of *A. callidryas* hatchlings from flooded and jiggled eggs across ages. Different letters indicate significant differences between ages from Tukey *post hoc* analyses of mixed models on each cue type respectively. Points indicate data from individual embryos (total *N*=87 hatchlings), jittered horizontally to improve visualization. Numbers show N of hatchlings measured per age, per cue type. Box plots show medians, IQR, and extent of data to ±1.5×IQR.

### Hatching latency

The latency to hatch (i.e., fully exit the egg) after stimulus onset varied with cue type and an age × cue type interaction (cue type: χ^2^=32.428, *P*=4.25e-07; interaction: χ^2^=12.819, *P*=0.0016), but there was no main effect of age (*P*>0.9). Across ages, latency was longer for flooded eggs (range: 2.29–21.41 min) than for jiggled ones (range: 0.25–3.58 min, Fig. 3A). Examining the effect of age within each cue type, hatching latency in response to hypoxia decreased by 73% from age 3–6 days (χ^2^=49.066, *P*=1.26e-10) and differed across all ages except 4 vs. 5 days (Tukey tests from GLMM, 4 vs. 5 d: *P=*0.173; 5 vs. 6 d: *P*=0.019; all others *P*<0.001). Across ages 4–6 days hatching latency in response to hypoxia decreased 53%, comparable to the 50% decrease in latency in response to physical disturbance (χ^2^=49.832, *P*=1.51e-11). Under physical disturbance, latency was similar at 4 and 5 days, but shorter at 6 days (Tukey tests from GLMM, 4 vs. 5 d: *P*=0.776; 5 vs. 6 d: *P*<1e-05; 4 vs. 6 d: *P*=0.012; Fig 3A). The variance in hatching latency was higher for flooded embryos than for jiggled ones at ages 5 and 6 days, but not at age 4 days (Fligner-Killeen tests, 4 d: χ^2^=2.50, *P*=0.114; 5 d: χ^2^=10.03, *P*=0.0015; 6 d: χ^2^=8.81, *P*=0.003; all df=1; Fig. 3A).

**Figure 3.**
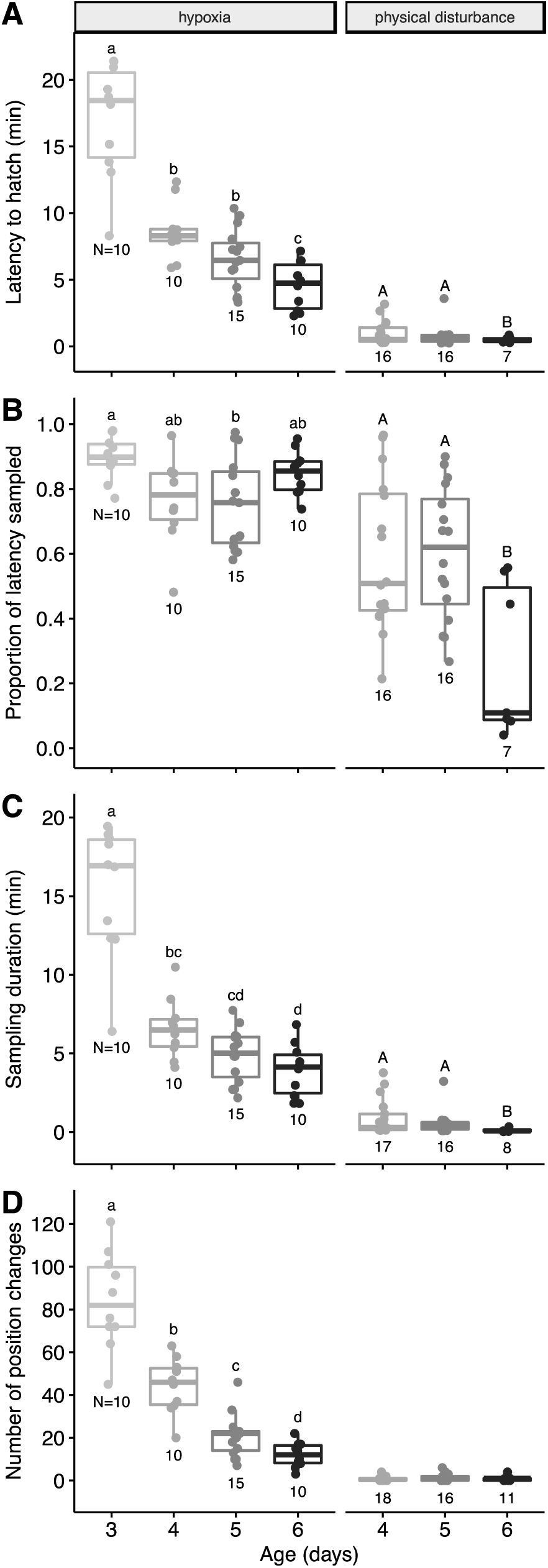
Ontogeny of latency to hatch and cue sampling by *A. callidryas* embryos in response to hypoxia and mechanosensory cues. (A) Entire latency period, from the onset of stimulation to complete exit from the egg, and (B) cue-sampling or risk assessment period, before hatching initiation, as a proportion of latency. (C) Duration of and (D) number of position changes during the sampling period. Different letters indicate significant differences between ages from Tukey *post hoc* analyses of mixed models on data from each cue type respectively. Numbers show N of individuals measured per age, per cue type. Data points are values from individual embryos. Box plots show medians, IQR, and extent of data to ±1.5×IQR.

### Information sampling and decision-making

The proportion of hatching latency spent in information-sampling and decision-making (i.e., the period from stimulus onset to hatching initiation) varied with age, cue type, and their interaction (gamma GLMM, age: χ^2^=21.282, *P*=0.0007; cue type: χ^2^=34.233, *P*=1.77e-07; interaction: χ^2^=18.854, *P*=8.05e-05). On average across ages, the sampling period accounted for a greater portion of the latency to hatching for flooded than for jiggled embryos (80.9 and 53.7%, respectively; Fig. 3B). Among flooded eggs, the main effect of age (χ^2^=10.502, *P*=0.015) was due to a difference between ages 3 and 5 days (Tukey test from GLMM, *P*=0.02; all other pairwise comparisons *P*>0.08, Fig. 3B). In jiggled eggs, the portion of latency spent sampling was smaller at age 6 days compared to both ages 4 and 5 days (main effect of age: χ^2^=10.338, *P*=0.0057, Tukey test from GLMM, 6 vs. 4 d: *P*=0.013; 6 vs. 5 d: *P*=0.0098, Fig. 3B).

Embryo age, cue type, and their interaction all had strong effects on the absolute sampling duration (gamma GLMM, age: χ^2^=42.717, *P*=4.22e-08; cue type: χ^2^=99.325, *P*<2.2e-16; interaction: χ^2^=26.234, *P*=2.01e-06; Fig. 3C) as well as on the number of position changes during cue sampling (Poisson GLMM, age: χ^2^=360.45, *P*<2.2e-16; cue type: χ^2^=149.04, *P*<2.2e-16; interaction: χ^2^=21.008, *P*=2.74e-05; Fig. 3D). Analyses within each cue type showed that sampling duration decreased with age in both flooded (χ^2^=49.716, *P*=9.18e-11) and jiggled eggs (χ^2^=14.913, *P*=0.0006).

The number of positions sampled also decreased with age in flooded eggs (χ^2^=363.07, *P*<2.2e-16, Fig. 3D) but not jiggled ones (*P*>0.1). Overall, jiggled embryos sampled for less time and assumed fewer positions compared to flooded ones (Fig. 3C,D). Even at age 6 days, flooded embryos changed positions on average 12 times (range: 3–22) while, across ages, 20 out of 45 jiggled embryos hatched without changing position and 15 more moved only once (range: 0–6 position changes). The mean duration of the sampling period and mean number of position changes at age 6 days were statistically greater than zero in both hypoxia (*t*-tests; duration: 3.92±1.69 min, *t*=7.3386, *P*=4.38e-05; position changes: 12.1±5.86, *t*=6.531, *P*=0.0001) and physical disturbance tests (duration: 0.11±0.11 min, *V*=36, *P*=0.0078; position changes: 0.91 ±1.14, *V*=28, *P*=0.015). Among flooded embryos, movement rates (i.e., the number of position changes per minute) decreased with age (χ^2^=29.596, *P*=1.68e-06) from 5.71±1.5 position changes per minute at age 3 days (range: 3.8–8.2) to 3.12±1.11 positions changes per minute at age 6 days (range: 1.0–4.9). Movement rates were specifically lower in flooded embryos at ages 5 and 6 days compared to ages 3–4 days (all *P*≤0.005).

### Hatching process

The entire hatching process, from hatching initiation to complete exit from the egg, took on average 55.8±66.3 s, across all ages and cue types (range: 0.09–6.11 min). We found significant effects of age, cue type, and their interaction on the duration of hatching, regardless of whether we excluded flooded eggs at age 3 days (gamma GLMM, age: χ^2^=18.245, *P*=0.0026; cue type: χ^2^=61.286, *P*=3.12e-13; interaction: χ^2^=10.191, *P*=0.0061). Across ages, the mean duration of the hatching process was shorter in jiggled than in flooded eggs (0.25±0.15 vs. 1.53±1.23 min, Fig. 4A). Among flooded embryos, those at age 6 days hatched the fastest, taking half as long, on average, compared to all other age groups (Tukey test from GLMM, all *P*<0.05, Fig. 4A). Conversely, in jiggled eggs, hatching took longer at age 6 days compared to ages 4 and 5 days (both P<0.0005, Fig. 4A). Thus, the overall fastest hatching (13.5±6.9 s) was for 4 and 5 d eggs in jiggling trials. The duration of hatching was more variable in response to flooding than jiggling at ages 4 and 5 days, but not age 6 days (Fligner-Killeen tests, 4 d: χ^2^=12.613, *P*=0.00038; 5 d: χ^2^=13.182, *P*=0.00028; 6 d: χ^2^=1.3331, *P*=0.25; all df=1; Fig. 4A).

**Figure 4.**
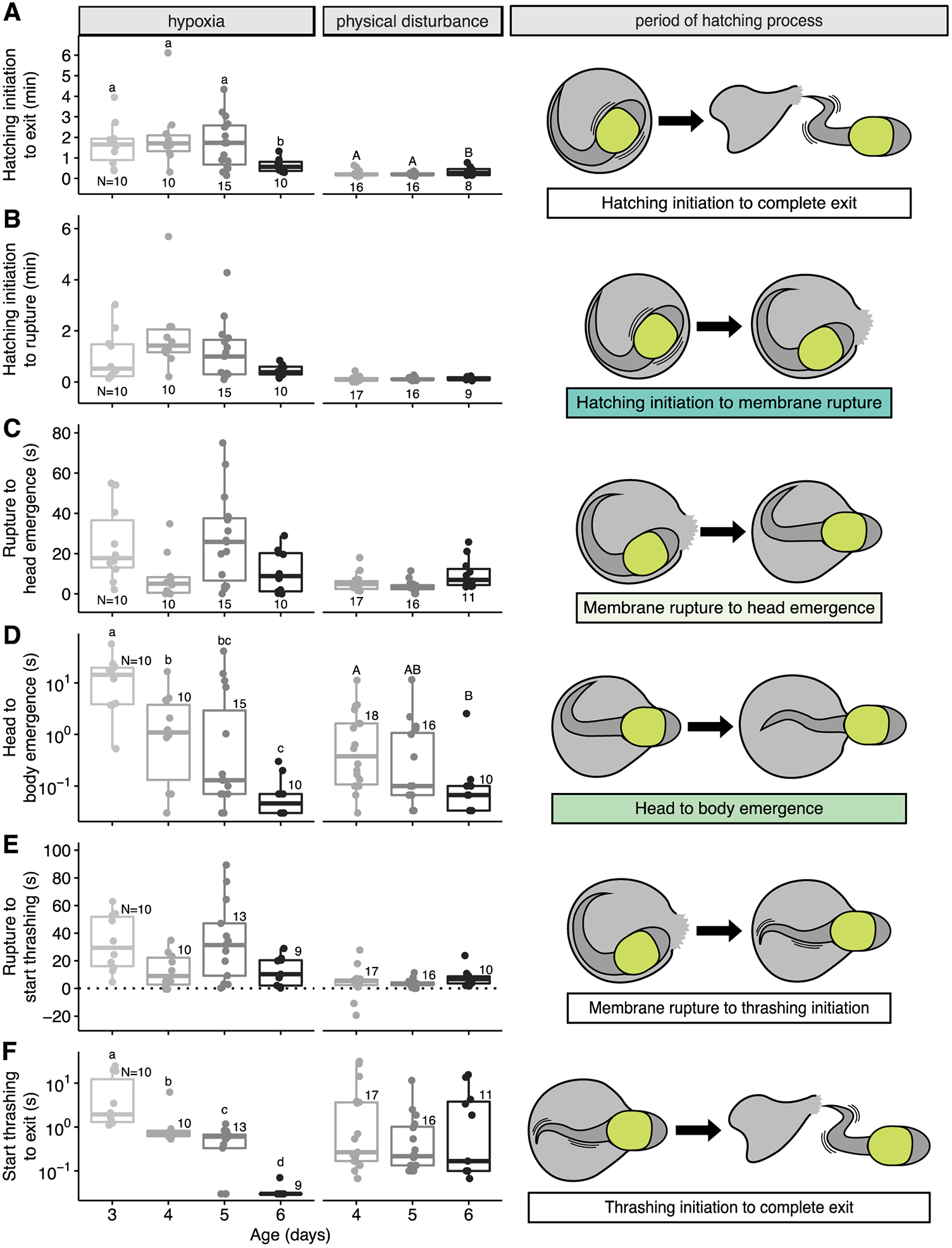
Ontogeny of hatching performance by *A. callidryas* embryos in response to hypoxia and mechanosensory cues. Periods from (A) hatching initiation to complete exit from the egg, (B) hatching initiation to membrane rupture, (C) membrane rupture to head emergence, (D) head to body emergence (excluding the tail), (E) membrane rupture to start of thrashing, and (F) start of thrashing to complete exit from the egg. Note log scale of Y-axis in D and F. Different letters indicate significant differences between ages from Tukey *post hoc* analyses of mixed models on data from each cue type respectively. Numbers show N of individuals measured per age, per cue type; some embryos did not thrash and hatching initiation was not recorded for all jiggled eggs. Points indicate data from individual embryos. Box plots show medians, IQR, and extent of data to ±1.5×IQR. Illustrations of periods within the hatching process (B–D) are color coded to match data presented in Figure 6.

Examining the period from hatching initiation to membrane rupture, we found a marginally significant effect of age (gamma GLMM, χ^2^=10.846, *P*=0.055) and no age × cue type interaction (χ^2^=1.1223, *P*=0.5705). In contrast, cue type strongly affected the initiation-to-rupture period (χ^2^=67.471, *P*<2.2e-16), which was shorter in response to jiggling (Fig. 4B). Because only shaking has been experimentally validated as a behavioral indicator of hatching enzyme release, we repeated this analysis on a dataset restricted to embryos that exhibited shaking; removing the eight cases with other hatching indicators weakened the age effect (χ^2^=9.9676, *P*=0.076). Variance in the initiation-to-rupture period was larger in flooding than jiggling at all ages (Fligner-Killeen tests, 4 d: χ^2^=16.08, *P*=6.09e-05; 5 d: χ^2^=15.98, *P*=6.4e-05; 6 d: χ^2^=5.24, *P*=0.022; all df=1; Fig. 4B).

The duration of the period from membrane rupture to head emergence varied with age, cue type, and their interaction (gamma GLMM, age: χ^2^=13.572, *P*=0.019; cue type: χ^2^=20.07, *P*=0.0002; interaction: χ^2^=8.6031, *P*=0.013; Fig 4C). This period was shorter for jiggled eggs than for flooded ones (6.0±5.3 s vs. 18.8±18.8 s, Fig. 4C). However, our analyses on each cue type separately found no effect of age on the duration of this period in either flooded or jiggled eggs (both *P*>0.1). The period from membrane rupture to head emergence was more variable in response to flooding than jiggling at ages 4 and 5 days, but not age 6 days (Fligner-Killeen tests, 4 d: χ^2^=4.15, *P*=0.042; 5 d: χ^2^=13.4, *P*=0.0003; 6 d: χ^2^=1.94, *P*=0.1633; all df=1; Fig. 4C).

We found no significant effect of cue type or age × cue type interaction on the period from head to body emergence (lognormal LMM, both *P*>0.5). Conversely, age was a strong predictor of the head-to-body-emergence period (χ^2^=42.075, *P*=5.68e-08). This result was evident in analyses of each cue type separately (hypoxia: χ^2^=29.65, *P*=1.64e-06; physical disturbance: χ^2^=7.07, *P*=0.03) and both showed a similar monotonic developmental decrease in the duration of this period (Fig. 4D). Variance in the period from head-to-body-emergence was not different across cue types at any age (Fligner-Killeen tests, all ages *P*>0.1, Fig. 4D).

The period from membrane rupture to the start of thrashing was affected by age, cue type, and their interaction (LMM, age: χ^2^=17.292, *P*=0.004; cue type: χ^2^=22.024, *P*=6.448e-05; interaction: χ^2^=6.921, *P*=0.0314; Fig. 4E). However, independent analyses within cue types found no effect of age in either flooded or jiggled eggs (both *P*>0.05). The rupture-to-thrashing period was shorter for jiggled eggs than for flooded ones (4.8±7.2 vs. 24.1±22.5 s). At age 4 days, a few embryos began thrashing prior to membrane rupture, resulting in negative values for this period (jiggling: –10.9 and –19.4 s; flooding: –0.58 and –0.59 s).

The period from membrane rupture to the start of thrashing was more variable in response to flooding than jiggling at ages 4 and 5 days, but not age 6 days (Fligner-Killeen tests, 4 d: χ^2^=4.41, *P*=0.036; 5 d: χ^2^=13.05, *P*=0.0003; 6 d: χ^2^=3.41, *P*=0.065; all df=1; Fig. 4E). The period from membrane rupture to head emergence depends strongly on the period from membrane rupture to thrashing onset (Gamma GLM, *R*^2^=0.57, χ^2^=172.41, *P*<2.2e-16).

Embryo age, cue type, and their interaction all had significant effects on the period from the start of thrashing to complete exit from the egg (lognormal LMM, age: χ^2^=39.458, *P*=1.92e-07; cue type: χ^2^=15.696, *P*=0.0013; interaction: χ^2^=11.515, *P*=0.0032; Fig. 4F). In independent analyses within cue type, we found the duration of this period decreased strongly with age in flooded eggs (χ^2^=53.448, *P*=1.471e-11) but did not vary with age in jiggled ones (*P*>0.5, Fig. 4E). Variance in the time from start of thrashing to exit was higher for jiggled embryos than for flooded ones at age 6 days but was not different at ages 4 or 5 days (Fligner-Killeen test, 4 d: χ^2^=3.15, *P*=0.076; 5 d: χ^2^=0.946, *P*=0.331; 6 d: χ^2^=10.758, *P*=0.001; all df=1).

Embryo age had a strong and significant effect on the incidence of body compression (binomial GLMM, χ^2^=29.265, df=5, *P*=2.057e-05, Fig. 5A), but neither cue type nor age × cue type interaction were significant (both *P*>0.1). At age 3 days, all embryos experienced body compression during hatching (Movie 3) while at age 6 days only one embryo from each cue type did; overall, the likelihood of body compression decreased an average of ∼30 and 12% per day in hypoxia and physical disturbance tests, respectively (Fig. 5A). The duration of body compression, for those embryos that experienced it, also appeared to decrease with age, but we found no significant age, cue type, or interaction effects on compression duration (gamma GLMM, all *P*>0.1, Fig. 5B).

**Figure 5.**
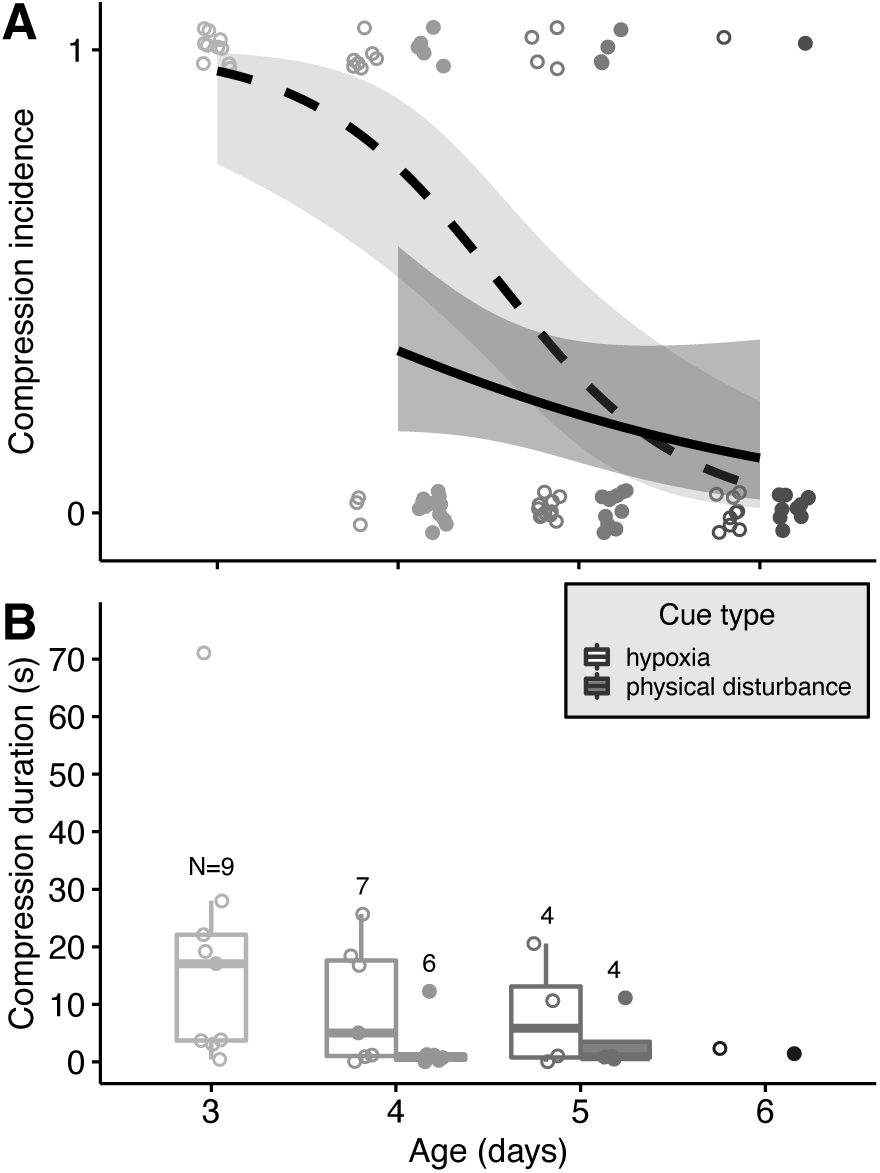
Ontogeny of body compression incidence and duration during hatching of *A. callidryas* embryos in response to hypoxia and mechanosensory cues. (A) Incidence of body compression during exit from the egg. Values of 1 and 0 indicate visible compression and lack thereof during hatching. Open (flooding) and closed (jiggling) data points are jittered horizontally and vertically to show data points within each age group and cue type. Dashed (flooding) and solid (jiggling) lines are predicted fits from binomial GLMMs; shading indicates the 95% confidence interval. (B) Duration of body compression during hatching. Points are jittered horizontally to show data within each age group and cue type. Open (flooding) and closed (jiggling) data points represent values for individual embryos and numbers show *N* of individuals measured per age, per cue type. Unfilled (flooding) and filled (jiggling) box plots show medians, IQR, and extent of data to ±1.5×IQR.

## DISCUSSION

In two common threat contexts, using cues in different sensory modalities, more developed embryos of *A. callidryas* assess risk and make hatching decisions based on shorter periods of cue sampling. Development also affects how well embryos execute their hatching decisions; some stages of hatching show a clear ontogenetic increase in performance, regardless of cue or context. However, context or cue type also affects both the process of risk assessment and hatching performance. Embryos spend less time in risk assessment during simulated attacks than they do under flooding. Moreover, some stages of hatching are consistently faster in simulated attacks than in flooding, suggesting that elements of hatching performance may reflect context-dependent variation in embryo effort more than developmental changes embryo abilities. Thus, adaptive context-and stage-dependent variation in embryo behavior appears to combine with information-and performance-related constraints to affect variation in risk-cued hatching.

### Information sampling and decision-making

Embryos of all ages showed a distinct cue-sampling period before beginning the hatching process; this was evident even at 6 days, when many embryos hatch spontaneously. Sampling duration decreased developmentally with both hypoxia and mechanosensory cues (Fig. 6), as predicted based on decreasing false-alarm costs. This extends findings from vibration playbacks at ages 5–6 d (Warkentin et al., 2019) to multiple threat contexts and a greater developmental range; consistently, more developed embryos base their hatching decisions on less information.

**Figure 6.**
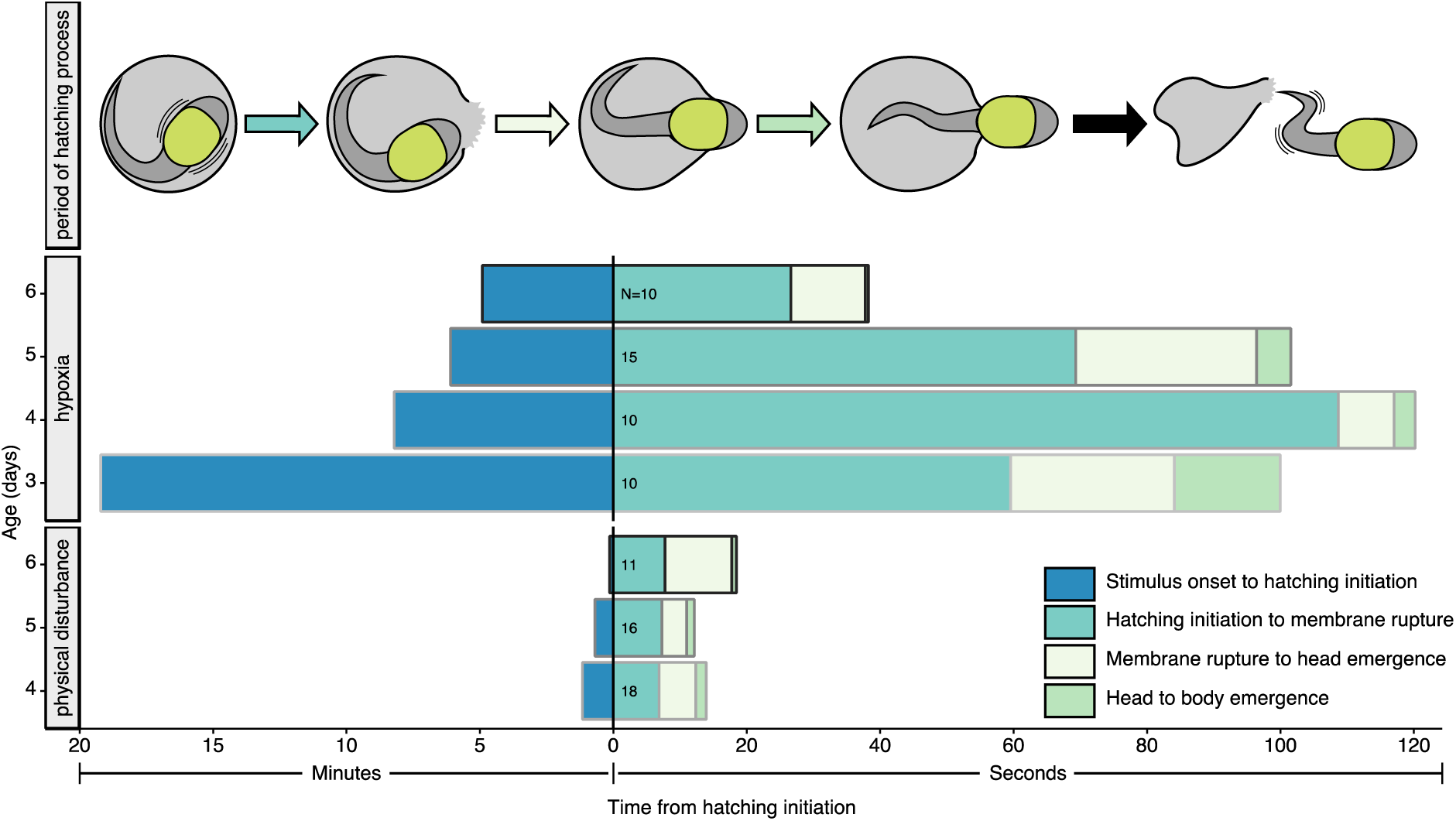
Mean durations of pre-hatching risk assessment and periods within the hatching process of *A. callidryas* embryos across ages and cue types. Note the different x-axis scales for periods before and after hatching initiation. Numbers show total N of individuals tested per age, per cue type; see Figs 3–4 for N of individuals measured per age, per cue type for each period. Periods within the hatching process are color coded to match data presented in Figure 4.

Cue type affected the sampling period even more than age; jiggled embryos initiated hatching 91% sooner after stimulus onset than flooded ones (Figs 3C, 6). This may reflect faster accumulation of both risk and information in predator attacks vs. flooding. First, egg-eating snakes can consume clutches in just a few minutes and, in snake attacks, the longer embryos spend assessing cues, the more their risk of mortality increases (Warkentin and Caldwell, 2009; Warkentin et al., 2007). Rapid mortality in predator attacks may have selected for rapid risk assessment and decision-making in response to mechanosensory cues. Conversely, even under strong hypoxia, submerged embryos survive and remain capable of hatching for over 20 min, enabling slower risk assessment (Fig. 3A). Under moderate hypoxia they may continue developing *in ovo* for days, accepting slower development to achieve a more advanced stage at hatching (Moskowitz et al., 2016; Snyder et al., 2018). Second, the rate and process of information acquisition differ between contexts. Embryos can sample mechanosensory cues passively, without moving. Vibration frequency spectra are immediately apparent (Caldwell et al., 2009; Warkentin and Caldwell, 2009) while temporal pattern information accrues continuously over time (Warkentin et al., 2007; Warkentin et al., 2019), even during periods of silence (Jung, 2021; Jung et al., *In press*). However, oxygen assessment is more active and time-consuming. Eggs in air contain hypoxic zones (Warkentin et al., 2005), so embryos must assess oxygen throughout their egg to determine that they are not simply facing the wrong way. This requires changing position, which stirs the perivitelline fluid, disrupting oxygen gradients (Rogge and Warkentin, 2008; Warkentin et al., 2005); embryos must then wait for the local oxygen level to stabilize to assess it. This slower process of information-gathering under hypoxia likely prolongs the risk-assessment period.

Position changes are a clear behavioral indicator of cue sampling by flooded embryos (Movie 1, Rogge and Warkentin, 2008). Both movement rate and the number of position changes before embryos initiate hatching decreased with age in hypoxia experiments suggesting that older embryos use less information for their hatching decision. The information embryos gain about oxygen per position might increase developmentally, for instance if the reach of the external gills or their oxygen-sensing capacity increases (e.g. number of neuroepithelial cells, Pan and Burggren, 2010); however, any such changes are probably small compared with the 86% decrease in positions sampled. In mechanosensory-cue experiments, only 56% of embryos changed position before hatching and only 22% moved more than once. Embryos have no need to move to sense these cues; indeed, self-generated motion might complicate the interpretation of lateral line and vestibular system input, hampering assessment of predation risk. Thus, embryos may avoid moving while assessing a physical disturbance. Rather than sampling, embryo position changes during egg-jiggling may reflect tactile-stimulated startle responses within the egg (Eidietis, 2006), or attempts to evade predators, since embryos are often directly bitten and poked during attacks (Hughey et al., 2015; Warkentin et al., 2006). Thus, while we consider position changes to be a useful indicator of sampling effort by *A. callidryas* during flooding, embryo movements during real or simulated predator attacks likely have other causes and may serve different functions.

### Latency to hatch as a proxy for sampling

The cue-sampling period of *A. callidryas* embryos represents a substantial, but variable, portion of latency to hatch (90–75% and 60–26% across ages, under flooding and jiggling, respectively; Fig. 3B), highlighting both the value and limitations of latency as an estimate of information sampling. Hatching latency provides more information than proportion hatched alone (Warkentin et al., 2019) and can reveal variation in behavior even when a stimulus induces all embryos to hatch (Jung et al., 2020). However, sampling represents a smaller fraction of latency with mechanosensory cues than hypoxia, particularly for older embryos. While latency is useful for comparisons within ages and cue types, and future studies of embryo risk assessment and hatching decisions should incorporate it along with hatching responses, more direct measurements of sampling will be essential in some contexts. The association between sampling periods and latency to execute behaviors also varies contextually in other taxa and at later life stages. For example, variation in sampling periods and behavioral performance both affect latency in mate choice by female túngara frogs. In phonotaxis experiments, females choose mates faster in response to complex advertisement calls and under higher light conditions (Bonachea and Ryan, 2011a; Rand and Ryan, 1981). While complex calls shorten the initial evaluation period and females leave the starting zone faster, higher light conditions result in faster movements towards the speaker. Females also seem to spend less time evaluating calls as simulated predation risk increases (Bonachea and Ryan, 2011b).

### Hatching process and performance

The entire hatching process, from initiation to complete exit, was faster in response to mechanosensory cues than hypoxia (15±9 vs. 92±74 s) and it was also less variable, particularly at ages 4–5 days (Figs 4A, 6). In previous work using a minimal mechanosensory stimulus, *A. callidryas* embryos at ages 5–6 days hatched in 21±11 s (range 6.5–49 s)(Cohen et al., 2016). In response to our stronger egg-jiggling stimulus, 6-day-old embryos performed similarly (21±13 s, 9–46 s) whereas 4–5-day-old embryos hatched even faster (14±7 s, 5.3–38 s, Fig. 4A). Faster—and more consistently fast—hatching in egg-jiggling vs. flooding experiments is congruent with our hypothesis that embryos perceiving an immediate threat of predation perform hatching behaviors closer to the limits of their ability. This may be especially so for younger embryos, that spend more time assessing risk (44±59 s at 4–5 d); the slightly slower, more variable hatching process of jiggled 6-d embryos follows very rapid risk assessment (6±6 s). Contributing to overall faster hatching, most periods within the process were shorter and less variable in mechanosensory-cue experiments (Fig 4A–C and E). The longer, more variable duration of hatching in flooded eggs seems likely to reflect individual behavioral decisions or effort, rather than the embryos’ full capability. This variation is consistent with substantial research on post-embryonic life stages, showing that animals’ realized performance often differs from their maximum capacity, with performance close to capacity only in a subset of contexts (Irschick and Garland Jr, 2001).

We anticipated that the transition from early to late HGC (age 3–4 days) and increase in late HGC abundance over development (Cohen et al., 2019) would improve embryos’ ability to rapidly rupture their vitelline membrane. Our data provide no evidence for such ontogenetic change (Figs 4B, 6). For jiggled embryos, the three shortest initiation-to-rupture times were at the youngest age (4 days, all <1 s). This period was longer in flooding, and more variable, with the two shortest times at age 3 days (8.6, 12.4 s) below any 4-day time (all ≥12.8 s). Thus, it appears this metric rarely reflects embryos’ full capacity, but comes closer under threat of predation. Most *A. callidryas* embryos use only a portion of their stored hatching enzyme per hatching attempt (Salazar-Nicholls et al., 2020), retaining enough to digest a second or even third escape hole if displaced (Salazar-Nicholls et al., 2017). Embryos might regulate enzyme release, using more to accelerate membrane rupture when seconds matter to escape predation or conserving it in case repeated hatching attempts are necessary. They might also behaviorally facilitate rupture by pressing their head against the membrane to increase HGC–membrane contact. Flooded embryos lose the oxygen gradient that helps them orient—and hatch—toward the exposed side of their egg, and in glass cups they also lose directional light cues; this increases the frequency of hatching complications and need for a second rupture site (Güell and Warkentin, 2018; Salazar-Nicholls et al., 2017). Moreover, in both snake and wasp attacks embryos sometimes move—or are pushed—away from their initial rupture site (KMW, observations from video recordings). Thus, the ability to make a second rupture might be an important element of hatching performance and/or developmental constraint (Salazar-Nicholls et al., 2017).

Like initiation-to-rupture, the period from rupture to head emergence, showed no age effect but was longer and more variable in flooded vs. jiggled embryos (Fig. 4C, E). This likely reflects behavioral decisions or effort, with flooded embryos using less of their capacity. Most embryos press their snout against the perivitelline membrane during, or soon after enzyme release and use thrashing movements to accelerate their exit (Cohen et al., 2016); indeed, the timing of thrashing onset explains much of the variation in head emergence. In contrast, the period from thrashing onset to exit showed different ontogenetic patterns across cue types. In flooding, this period decreased developmentally, whereas jiggled embryos showed intermediate values with no developmental change (Fig. 4F). This may reflect a combination of effort and constraint. The earlier onset of thrashing in jiggling experiments suggests that embryos perceiving predation risk attempt to behaviorally hasten their exit; however, later thrashing of flooded embryos could allow time for greater membrane digestion. Consistent with this, in flooding most 6-day and several 5-day embryos exited with a single tail flick, in <0.03 s, whereas jiggled embryos were never that fast. Moreover, the long thrashing-to-exit periods of the youngest hatching-competent embryos, even after a post-rupture wait, suggest a stronger developmental constraint.

Both the period from head to body emergence and incidence of body compression during this period decreased strongly with age, with no cue effect, indicating developmental constraints (Figs 4D, 5, 6). Several morphological changes seem likely to improve these elements of hatching performance. Over the plastic hatching period, embryos grow longer, more muscular tails and become more streamlined as yolk is transformed into other tissues (Warkentin, 1999b). At age 3 days, embryos have small heads and their yolk-filled bellies are the widest part of their body (Fig. 1B, Movies 1, 3); even with active thrashing, their exit typically slows once their head has emerged and body compression is always evident as their bulbous yolk squeezes through the rupture site (Fig. 5). In contrast, by age 6 days embryos’ heads are wider than their bodies, facilitating a quick, smooth exit once the head has emerged (Movie 2). This acceleration of emergence and lower incidence of compression also suggests that older embryos make larger holes in the membrane; they might release more hatching enzyme and their broader heads could enable enzyme delivery to a larger area of membrane (Salazar-Nicholls et al., 2020). Moreover, greater reach, propulsive area, and strength may increase thrashing effectiveness and exit speed as embryos develop.

### Understanding variation in latency to hatch

The fact that *A. callidryas* use multiple cue types to hatch in multiple risk contexts enables consistent developmental changes in the process, due to ontogenetic adaptations or release from developmental constraints, to be distinguished from context-specific differences in behavior or performance, which may also reflect adaptive variation or environmental constraints. More generally, distinguishing the component processes that comprise cued hatching, and when each occurs, can facilitate identification of factors and mechanisms that generate variation at each stage of the process (Fig. 7). This framework for assessing determinants of variation in hatching should be applicable within and among species.

**Figure 7.**
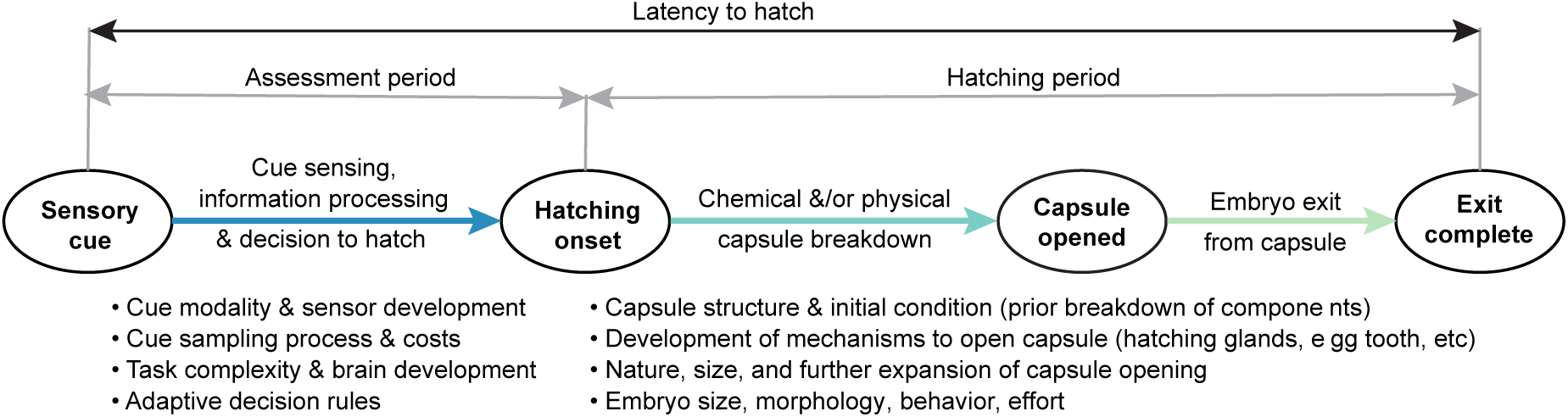
Elements of variation in hatching latency. Cue-to-exit time includes both assessment and hatching periods (grey), which may be further subdivided, as illustrated for hatching. The processes within each period (colored arrows) depend on multiple factors that vary with evolution, development, and environmental context (examples listed).

Published measures of cued hatching timing rarely distinguish assessment from hatching periods, and hatching may also be initiated by internal, developmental events rather than external cues. However, all embryos must hatch from their protective capsules and both the overall duration of the process and the dominance of stages within it vary. When hatchlings of low mobility remain on or near their egg clutch, fitness trade-offs at hatching and selection for hatching speed may be limited. Moreover, the capsule structure and mechanism for opening it may impose constraints. Some aquatic amphibian eggs hatch very slowly, with dissolution or emergence from jelly coats, enzymatic degradation of the entire vitelline membrane, then further localized weakening, rupture, and emergence taking about half the embryonic period (e.g., Carroll Jr and Hedrick, 1974; reviewed in Cohen et al., 2016; Yoshizaki, 1978). Many reptiles and birds are also slow to hatch, taking many hours to days to completely emerge from the egg after pipping (Doody, 2011; Oppenheim, 1972; Oppenheim, 1973; Visschedijk, 1968). At the other extreme, when rapid, precisely timed emergence is essential for hatchlings to exploit a transient and essential opportunity, or escape from a sudden threat, hatching latency may be very brief. The Californian grunion, *Leuresthes tenuis*, a terrestrially incubated fish that hatches in response to agitation by waves, emerges <1.5 min after stimulus onset (Speer-Blank and Martin, 2004), while delicate skinks hatch in <10 s in response to a simulated predator attack (Doody and Paull, 2013). Moreover, some parasitic flatworms (Monogenea) hatch just 2–4 s after exposure to skin mucus from their host fish, physical disturbance, or sudden shadows (reviewed in Whittington and Kearn, 2011). Although we do not know their separate durations, such fast responses must involve both a rapid assessment and decision process and speedy mechanisms to rupture and exit the capsule.

The duration of the hatching period depends on egg capsule structure and the mechanisms used to open and exit from it (Fig. 7). Many species, including flatworms (Kearn et al., 1999), nematodes (Mkandawire et al., 2022), insects (Donoughe, 2021), and gastropods (Rawlings, 1999) have opercula or polar plugs that facilitate rupture and provide predefined exit sites. Conversely, bird embryos must create an exit site by tearing internal membranes and cracking their eggshell bit by bit around an arc or circle (Hamburger and Oppenheim, 1967; Oppenheim, 1972; Oppenheim, 1973). In many aquatic-breeding anurans the entire vitelline membrane is slowly digested by gradual hatching enzyme release, while *A. callidryas* use rapid, localized enzyme release to digest a small escape hole (Cohen et al., 2018; Yamasaki et al., 1990; Yoshizaki, 1978; Yoshizaki and Katagiri, 1975). Anticipatory changes to the capsule, prior to the hatching decision, can also speed hatching. For instance, while embryos of some monogenean flatworms enzymatically soften their opercular cement on perceiving a cue, hatching in 4–5 min, others pre-weaken the cement and can exit in as little as 2 s (Whittington and Kearn, 2011). Hatching may also be linked to, and slowed by, associated developmental processes. For instance, in reptiles, complete emergence after pipping can be delayed for days as the hatchlings wait for their yolk sac to be internalized (Pezaro et al., 2013). Alternatively, to emerge rapidly reptiles may sacrifice energy reserves, leaving yolk behind when they exit (Doody and Paull, 2013).

Cued hatching requires mechanisms linking environmental context to hatching timing, thus cue sensing is often the first step in the process. In the simplest case, if embryos hatch in a single context, using a consistent, distinctive cue, sensing the cue could simply trigger a reflexive hatching response. This might occur, for instance, in the extremely rapid responses of some monogenean flatworms to host mucus or sudden darkness (Whittington and Kearn, 2011). A simple circuit from transient hindbrain photoreceptors to hatching gland cells mediates light-inhibited/dark-induced hatching in Atlantic halibut (Eilertsen et al., 2018). In other cases, if embryos use multiple cue types, information accrues more slowly, and hatching cues are less distinct from the background, information processing will be more complex and the assessment period longer (Jung et al., *In press*; Warkentin and Caldwell, 2009). In such cases, as with behavioral decisions at later life stages, assessment or cue-sampling costs may also affect the decision process and timing (Warkentin et al., 2007). Moreover, developmental changes in the costs of missed cues or false alarms may select for ontogenetic adaptations in assessment and decision strategies (Jung et al., 2021; Warkentin et al., 2019).

Hatching is an essential, irreversible behavior that causes greater physiological and ecological change than most animal actions. Across taxa, embryos gather information from outside their egg to make informed hatching decisions that improve their immediate and ultimate survival (Du and Shine, 2022; Warkentin, 2011). The rapidly developing cognitive and physical capabilities of embryos make hatching an excellent, and underutilized, process in which to study how development affects information use and behavioral performance across contexts.

## Acknowledgements

We thank Alina Chaiyasarikul, Karina Escobar, Su Jin Kim, Nora Moscowitz, Sonia Pérez Arias, María José Salazar Nicholls, Crystal Tippett, and Angelly Vasquez for assistance with egg collection and care, and Richard Wassersug for discussions of embryo cranial kinematics. We thank members of the Gamboa Frog Group at STRI and Egg Science Research Group at BU for discussions of this work at multiple stages, and Ben Johnson for comments on the manuscript

## Author contributions

Conceptualization: K.M.W.; Methodology: B.A.G., J.J., J.C., A.A., K.M.W.; Formal analysis: B.A.G.; Investigation: B.A.G., J.J., J.C., A.A., K.M.W.; Resources: K.M.W.; Data curation: B.A.G.; Writing – original draft: B.A.G.; Writing – editing: B.A.G., K.M.W.; Writing – review: B.A.G., J.J., J.C., A.A., K.M.W.; Visualization: B.A.G., K.M.W.; Supervision: K.M.W.; Project administration: K.M.W.; Funding acquisition: K.M.W.

## Ethics approval

Research was conducted under permits from the Panamanian Ministry of the Environment (SC/A-15-14, SE/A-46-15, SE/A-59-16), STRI IACUC protocol 2014-0601-2017, and BU IACUC protocol 14-008.

## Data availability

Data are available from the Dryad Digital Repository (details to be added)

## Supplementary Information

**Movie 1. Flooding and position changes or oxygen-sampling behavior**. The clip shows an *A. callidryas* embryo at 3 days inside a custom-made glass cup used in hypoxia-cued hatching tests, as it is submerged under boiled, degassed water and changes positions. The video was recorded using a Canon DSLR and MPE-65 mm macro lens and plays first in real time and then at 5x speed.

**Movie 2. Jerking behavior, buccal cavity compression, membrane rupture, and exit from the egg**. *A. callidryas* embryos 6 days of age display jerking behavior and a buccal cavity compression while flooded. The first embryo moves quickly, but does not change position, using a strong and abrupt axial muscle contraction (jerk). The second embryo moves its lower jaw, briefly changing the shape of its snout, but does not gape open its mouth or jerk its body (buccal cavity compression). This behavior is followed by rupture of the egg membrane and exit from the egg capsule. The videos were recorded using a Canon DSLR and MPE-65 mm macro lens and play in real time.

**Movie 3. Body compression, thrashing, and exit from the egg**. An embryo 3 days of age shows extensive body compression and thrashing effort as it exits from the egg capsule during flooding. The video was recorded using a Canon DSLR and MPE-65 mm macro lens and plays in real time.

